# BlobToolKit – Interactive quality assessment of genome assemblies

**DOI:** 10.1101/844852

**Authors:** Richard Challis, Edward Richards, Jeena Rajan, Guy Cochrane, Mark Blaxter

**Author notes:** Corresponding author: Richard Challis, Wellcome Sanger Institute, Cambridge CB10 1SA, UK.

## Abstract

Reconstruction of target genomes from sequence data produced by instruments that are agnostic as to the species-of-origin may be confounded by contaminant DNA. Whether introduced during sample processing or through co-extraction alongside the target DNA, if insufficient care is taken during the assembly process, the final assembled genome may be a mixture of data from several species. Such assemblies can confound sequence-based biological inference and, when deposited in public databases, may be included in downstream analyses by users unaware of underlying problems.

We present BlobToolKit, a software suite to aid researchers in identifying and isolating non-target data in draft and publicly available genome assemblies. BlobToolKit can be used to process assembly, read and analysis files for fully reproducible interactive exploration in the browser-based Viewer. BlobToolKit can be used during assembly to filter non-target DNA, helping researchers produce assemblies with high biological credibility.

We have been running an automated BlobToolKit pipeline on eukaryotic assemblies publicly available in the International Nucleotide Sequence Data Collaboration and are making the results available through a public instance of the Viewer at https://blobtoolkit.genomehubs.org/view. We aim to complete analysis of all publicly available genomes and then maintain currency with the flow of new genomes. We have worked to embed these views into the presentation of genome assemblies at the European Nucleotide Archive, providing an indication of assembly quality alongside the public record with links out to allow full exploration in the Viewer.

## Introduction

Genome sequences are part of the basic data economy of modern bioscience. Using assembled genomes, it is possible to identify loci underpinning key traits of interest, discover the regulatory logic of gene expression, investigate disease processes, and explore the evolutionary histories of genes and species. These research programmes rely implicitly on the correctness of the genome sequences. Errors in genome sequences risk distracting or even derailing their effective use.

Assembly of true genome sequences from reads shorter than the length of a replicon remains a difficult task (Ekblom and Wolf 2014). This task is made more complex when isolation of the original samples or the processing of DNA to generate the raw sequence data cannot avoid contamination of the target genome with DNA from non-target sources (Salter et al. 2014; Salzberg et al. 2005). Sequencing instruments are agnostic as to species-of-origin of the fragments they are tasked with processing, and thus a contaminated sample will result in a contaminated raw dataset. If insufficient care is taken during the assembly process, this can mean that the final assembled genome is a mixture of data from several species, and cannot be used as a good representation of the target species (Merchant, Wood, and Salzberg 2014). Downstream, this can result in erroneous attribution of biochemical or genetic properties to the target species that are actually derived from the contaminants’ genomes (Artamonova et al. 2015; Arakawa 2016).

However, not all “contaminants” are uninteresting. Many eukaryotic species live in close biological association with symbionts, and many bacteria exist in, and can only be grown as, consortia of interacting species (López-García, Eme, and Moreira 2017). In these systems genome sequencing aims to reconstruct the genomes of all the independent species and strains involved (Kumar and Blaxter 2011).

We are developing BlobToolKit, a software suite that will aid researchers in identifying contamination before it is erroneously blessed as being part of a target genome and to separate sequences that belong to different members of biological consortia. BlobToolKit is based on BlobTools written by Dominik Laetsch (Laetsch and Blaxter 2017) which was in turn based on the original Blobology pipeline from Sujai Kumar (Kumar et al. 2013). We present a toolkit that has been rewritten in its entirety to make use of advanced web frameworks and visualisation. Like its progenitors, BlobToolKit uses GC proportion and coverage as two major axes on which contigs or scaffolds from an assembly can be displayed.

GC proportion is consistent within most genomes, with a distribution around a mean value. Genomes can have regions of differing composition yielding bi-(or multi-) modal distributions. Genomes from different taxa present in a mixed sample frequently have different GC proportion, permitting a primary separation on this axis. The read coverage of each contig or scaffold in an assembly is an estimate of the relative stoichiometry of the replicon from which it derives. All the contigs or scaffolds from one species should have the same coverage, barring the presence of organelles (usually high coverage relative to the nuclear genome), sex chromosomes (50% coverage in the heterogametic sex) and uncollapsed haploid segments (again 50% coverage). Contaminant or cobiont genomes will have different, internally consistent stoichiometry and thus can be distinguished on this axis.

To provide initial identification of sets of contigs or scaffolds from distinct taxa, BlobToolKit also decorates each scaffold or contig with a taxonomic attribution based on similarity to sequences in reference databases, as assessed by BLAST (Altschul 1997) or Diamond (Buchfink, Xie, and Huson 2015). This taxonomic attribution is tentative due to the presence of mis-annotated records in the public databases. In conjunction with GC proportion and coverage measures this serves to highlight clusters (or blobs) of contigs that share distinct properties and coherent taxonomic source.

This richly marked-up annotation of the assembly makes it possible to assess whether it derives from single or multiple source organisms. The BlobToolKit data can be used to separate contigs and scaffolds (and the reads that generated them) into separate bins for subsequent reanalyses. BlobToolKit can be used as part of the process of genome assembly, playing a role both in separating raw input data for assembly of distinct components and in quality assurance of the final product. For genome assemblies released publicly, BlobToolKit can be used to provide quality assurance and to identify issues that should be taken into consideration in downstream reuse of the data.

Here we present the latest version of BlobToolKit, show how it can be used to probe the integrity of genome assemblies, describe the visualisations available and present snapshots of our ongoing BlobToolKit analyses of all eukaryotic genome assemblies available in the European Nucleotide Archive (ENA) (Amid et al. 2019).

## BlobToolKit

All BlobToolKit code is freely available under open source licenses from https://github.com/blobtoolkit. Distinct components are placed in four repositories: **BlobTools2** (command line tools to create and filter datasets), **Specification** (a formal specification and validator for the JSON-based data format), **Viewer** (interactive dataset visualisation), and **INSDC-pipeline** (a Snakemake pipeline to run the BlobToolKit workflow on publicly available datasets).

### BlobTools2

**BlobTools2** is a command line program to import a genome assembly together with BLAST, Diamond, read mapping and BUSCO analysis output files to generate a dataset that can be filtered using the command line and/or explored interactively in a web browser using the BlobToolKit **Viewer**.

#### *BlobDir* format

**BlobTools2** is a re-implementation of BlobTools (Laetsch and Blaxter 2017), written in python3 and based around a *BlobDir* directory of JSON format files. This data structure has been chosen as it can be easily validated using JSON-schema and is highly extensible. Separate JSON files contain distinct attributes of the assembly, with one entry per contig or scaffold. The attributes include GC proportion, length, coverage from a single sequencing library, taxonomic inference based on BLAST hits. Because the attributes are treated as generic datatypes (identifiers, variables, categories, arrays of categories or variables and arrays of arrays), it is possible to incorporate results from new analyses without making significant changes to the codebase. Field metadata are collated in a single JSON file allowing basic dataset information to be accessed without loading the full set of values. JSON is the native format for the JavaScript-based BlobToolKit **Viewer** and the typical patterns of use require computation across all data for a given attribute at once. Because the **Viewer** architecture inverts the usual server-client model, pushing computation to the client, this *BlobDir* format was favoured for efficiency of data access over alternatives such as SQLite or HDF5.

#### Adding data to a *BlobDir*

##### Assembly

The minimum input required to create a new *BlobDir* dataset is a FASTA format assembly sequence file. This is parsed to generate a list of sequence identifiers, along with a set of basic, per-sequence statistics (length, GC proportion and undefined [N] bases). Additional metadata, including assembly accessions and taxonomic information can be provided for inclusion in the dataset metadata and, if an NCBI taxonomy ID (taxid) is provided, expanded taxonomic lineage details will be included. The *BlobDir* can be modified, for example, to add attributes based on new analyses, using the **BlobTools2** *add* command.

##### Coverage

Both base and read coverage are calculated for each contig by parsing read alignment files in BAM, SAM or CRAM formats using the pysam library (https://github.com/pysam-developers/pysam).

##### Taxonomy

Taxonomy information is assigned to contigs and scaffolds through parsing of similarity searches of taxonomically-annotated sequence databases. Rather than simply use a single, top-scoring hit for each contig or scaffold, **BlobTools2** uses simple taxonomy rules (taxrules) to deliver a best-supported assignment. **BlobTools2** deploys taxrules introduced in BlobTools to assign putative taxonomic associations to sequence contigs: *bestsum* (total bitscore of all hits across all databases) and *bestsumorder* (total bitscore from a single database search, with scores taken from successive databases for contigs or scaffolds that failed to identify hits in the first). In a typical use case a file of NCBI BLAST+ *blastn* hits to the NCBI nt nucleotide database and a file of Diamond blastx hits to the UniProt/SwissProt database are supplied to be processed under one of these taxrules to generate a set of JSON files. For each of eight taxonomic ranks from superkingdom to species, files are generated containing the most likely taxon name, the summed bitscore of all hits to that taxon, a c-index value indicating the number of alternate taxa at that rank, and taxon names for every hit to each contig or scaffold. An additional file shows the location, score and taxid for every hit, information that is independent of the taxonomic rank under consideration. Results are split across multiple files to allow faster access to individual components during subsequent analyses.

#### BUSCO

As an example of the incorporation of new analyses, BUSCO (Benchmarking Universal Single-Copy Orthologues), a widely used tool for quality assessment of genome assemblies (Waterhouse et al. 2017) generates a sparse annotation where a few contigs are decorated with the presence of a BUSCO reference gene. **BlobTools2** incorporates BUSCO using the same basic datatype as BLAST hit distributions. The only unique code occurs in a specially written parser module for the BUSCO file format.

##### Hyperlinks

Hyperlink templates can also be added to the *BlobDir* metadata to allow hyperlinks from assembly/taxon identifiers, individual sequence identifiers or individual BLAST/Diamond hits to external resources.

#### Applying filters

**BlobTools2** supports filtering of assembly files, read files and of *BlobDir* datasets based on values of any of the constituent attributes. Variable attributes support filtering based on maximum and/or minimum values, category attributes may be filtered by presence or absence of one or more keys and individual records can be filtered with lists of identifiers to keep or exclude. Filtering of input files can assist in the process of iterative assembly improvement, while filtering of datasets may allow more detailed interrogation of subsets of the data in the BlobToolKit viewer without the need to repeat analyses or filter analysis outputs for re-importing.

### Specification

The BlobToolKit **Specification** describes the file formats required by **BlobTools2** and the BlobToolKit **Viewer** and includes a validator that tests a *BlobDir* dataset for departures from the specification. Use of JSON format allows validation with JSON-schema. While basic validation is possible with a static schema, the validator generates and tests against dynamically generated schemas to allow for the dependence of some metadata values on the presence and content of data in field-specific files. Validation includes type checking, testing for presence and content of expected files and assessing metadata ranges against the values present in corresponding field files.

### BlobToolKit Viewer

The BlobToolKit **Viewer** allows interactive exploration of *BlobDir* datasets produced by **BlobTools2**.

#### Application programming interface

All data in a *BlobDir* can be made available through an application programming interface (API) implemented using the Express Node.js web framework (https://expressjs.com/). The API provides search functionality against entries in the *assembly* and *taxon* sections of the metadata along with direct access to datasets, fields and individual records within fields. Full API documentation is available at https://blobtoolkit.genomehubs.org/api-docs/.

#### Interactive data exploration

The BlobToolKit **Viewer** presents data retrieved via the API in a set of interactive views for dataset visualisation, exploration and filtering. The **Viewer** is built on the React (https://reactjs.org) JavaScript library. It makes extensive use of Redux and reselect frameworks to allow real-time interaction with genome-scale datasets in client web browsers. This makes it practical to host large numbers of publicly accessible datasets on a server with a relatively small footprint. For datasets that are too large to be processed on the fly (including those with millions of contigs), pre-generated static image files can be served in place of the interactive views. Interactive plots are powered by the d3 data visualisation library (https://d3js.org) and all plots can be exported directly as PNG or SVG image files.

##### Filters view

The **Viewer** supports the same set of filter parameters as **BlobTools2**. Filter controls provide a graphic representation of category or variable distributions. To reduce network overheads, only data for active fields are loaded into the browser and the filter view provides an indication of which data are currently available. All views update instantly based on changes to filters.

##### Blob view

Blobology and BlobTools introduced the blob plot with contigs represented as circles, with areas proportional to contig length. This representation has several computational and interpretation issues. Circles are computationally expensive to plot, and rendering of datasets with many contigs (some published assemblies have over 1 million) makes it impossible to see all the data. While the scaled circle view is available in the **Viewer**, the default is to bin data into squares or hexagons of GC proportion-coverage space. In addition to resolving the problems with circles identified above, binning makes it possible to interactively select contigs within a chosen GC proportion-coverage bin. A square-binned blob plot of GC proportion *vs*. coverage is the default view when opening a new dataset with the **Viewer**. The squares are scaled to the square-root of the sum of lengths of contigs within each bin and coloured by best-matching phylum.

If used with a single scaling function and parameter set, binning has disadvantages, especially where the reduced prominence of minor sets of contigs can make it harder to identify cobionts. To overcome this, several scaling parameters can be modified, and viewing the plot while changing these parameters can be a useful way to explore features that are not immediately clear in a static image. The resolution of the plot can be adjusted, making the bins larger to facilitate the selection of major features or smaller to highlight fine-scale patterns, such as the off-axis bimodality associated with heterozygosity. The default square-root scaling can be adjusted to log or linear scales to increase or reduce the prominence of smaller values. The reducer function used to convert the values for each contig into a single value can also be adjusted from the default (sum) to show the minimum, maximum, mean or count of values in each bin.

All of these options are available for plots of any variable in the dataset against any other variable, for example, to allow coverage *vs*. coverage plots to identify contigs that are only supported by one sequencing library. Categories may be assigned based on any of the taxonomic ranks that have been calculated.

##### Cumulative view

The Cumulative view is a commonly used representation of the fraction of the genome that is represented as size-ordered contigs are added to the assembly. These plots also show this cumulative distribution broken down according to taxonomic assignments (the default is by phylum) and allow these separate curves to be stacked to show cumulative span by taxon.

##### BUSCO view

If BUSCO scores are added to a dataset, the BUSCO view shows a summary of the counts in each BUSCO category (complete, fragmented, etc.) under the current set of filters. It also allows selection of all contigs within a BUSCO category so that their distribution can be seen in the Blob view or the contigs can be inspected in the Table view. These interactions with other views make it possible to assess the impact of possible cobionts on the overall BUSCO score for an assembly.

##### Snail view

The Snail view is a reimplementation of interactive assembly statistic plots introduced in the Lepbase project (Challis et al. 2016). These capture a rich variety of assembly properties in a single dynamic graphic. Snail plots can highlight specific features of an assembly that may not be immediately apparent from tabulated data.

##### Table view

The Table view shows information for each contig for each currently active attribute. The available columns can be controlled by activating or deactivating individual attributes in the Filters view. The default columns show the GC proportion, length, coverage and taxonomic assignment that are used to generate plots in the Blob and Cumulative views. Individual records can be selected (either to view their position in the Blob view or for use in filtering) and rows can be sorted according to selected status or by any of the attribute values.

##### Hit view

The Hit view shows the distribution of sequence similarity hits to sequence databases along a single contig and can be accessed from the Table view of contigs, and is particularly useful for investigating contigs or scaffolds with unexpected or conflicting taxonomic attribution. The hyperlink functionality can be used to embed links to associated records in public sequence databases.

##### Detail view

A subset of dataset metadata is presented in a tabular Detail view, together with optional links to external resources. Full dataset metadata can be retrieved in JSON format.

##### Reproducible analyses

Sharing analyses reproducibly is critical, particularly when many choices have been made to generate a particular filtered dataset or image. To aid in reproducibility the **Viewer** encodes query parameters within the URL for the displayed data. Parameters developed during interactive filtering can be applied in **BlobTools2** (specified individually or using the entire URL or query string) to filter input files and *BlobDir* datasets. Selection-based filters are not stored in the URL due to the potential number of identifiers involved. Selections can be exported and imported *via* a List menu, which will export a JSON format file that includes a complete list of identifiers based on the current filters, including selections. This file also contains a summary of URL parameters and filtered dataset statistics (including BUSCO scores, span and N50 by taxon, etc.) and can be used to specify filter parameters used within **BlobTools2**.

##### Access to views

**BlobTools2** provides a *view* command that uses the Selenium WebDriver to provide non-interactive access to all plot types. For datasets with millions of contigs that are too large for practical interactive exploration, use of *view* provides a way to generate static images that will not display in the interactive mode.

### INSDC-pipeline

**INSDC-pipeline** is a reusable Snakemake (Köster and Rahmann 2018) pipeline to run analyses on publicly available, International Nucleotide Sequence Database Collaboration (INSDC; http://www.insdc.org/) public eukaryotic genome assemblies. We built the pipeline to automate the generation of *BlobDir* datasets from the available data, including retrieval and formatting of database files, retrieval of sequences for each assembly and the associated raw read files, read mapping, BLAST and Diamond searches, and BUSCO analyses (Figure 2). We have made the results available on a public instance of the BlobToolKit **Viewer** at https://blobtoolkit.genomehubs.org/view (Table 1).

**Table 1.** Summary of assemblies analysed and available* at https://blobtoolkit.genomehubs.org/view on 13th November 2019.

| *Kingdom* | *Species* | *Assemblies* |  |  |
| --- | --- | --- | --- | --- |
|  | *Total* | *Total* | *With reads* | *Without reads* |
| **Fungi** | 830/2016 | 1689/4992 | 1211/2787 | 478/2205 |
| **Metazoa** | 570/1437 | 743/2160 | 376/1311 | 367/849 |
| **Viridiplantae** | 180/545 | 232/839 | 121/439 | 111/400 |
| **Other Eukaryota** | 275/353 | 516/768 | 204/398 | 312/370 |
| **Total** | **1855/4351** | **3180/8759** | **1912/4935** | **1268/3824** |

\*For each kingdom within Eukaryota, the numbers of assemblies analysed/available are shown. Values were obtained through using a scripted query of the ENA and BlobToolKit APIs described in File S1.

**Figure 1.**
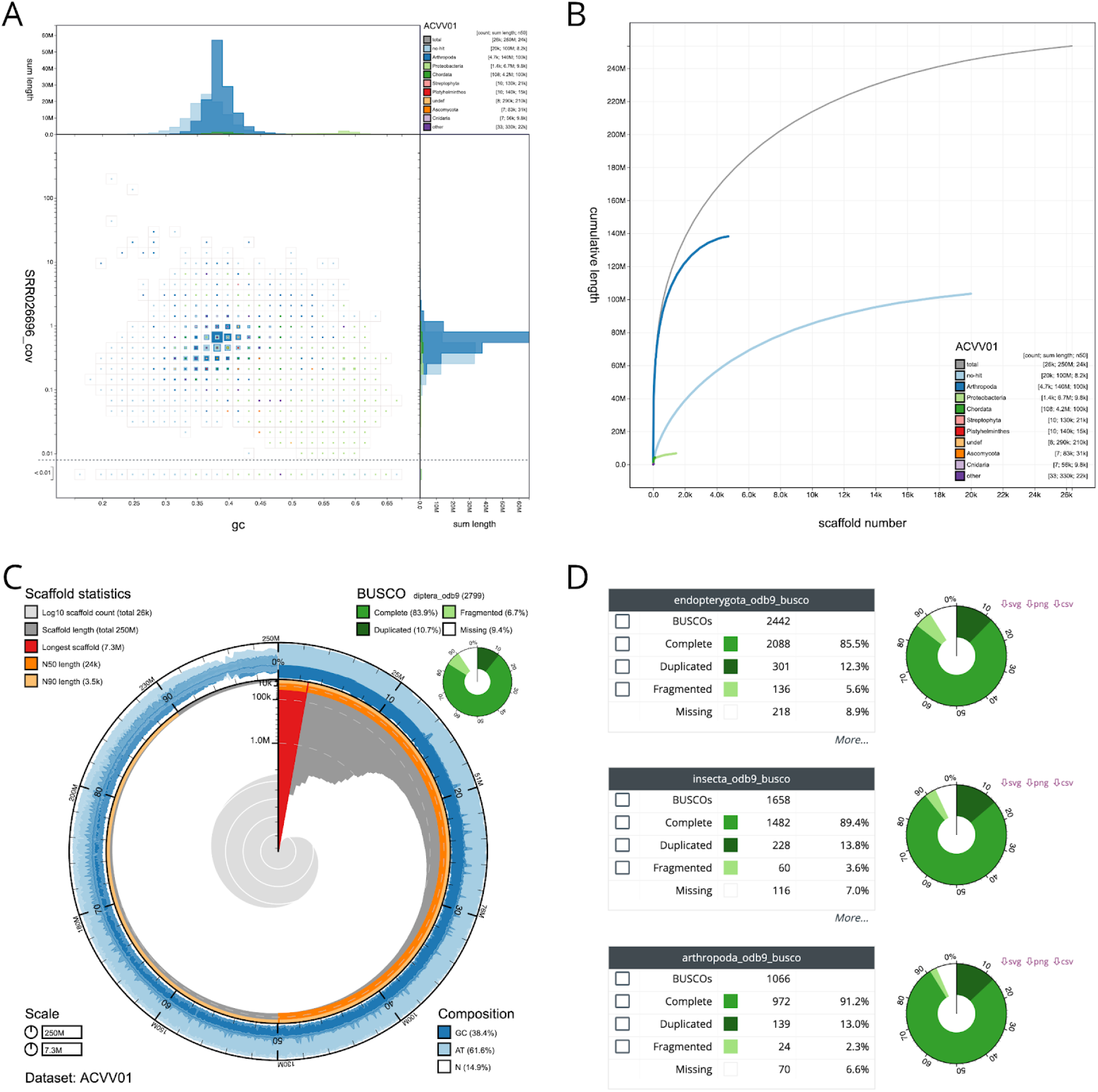
Assembly views available in the BlobToolKit Viewer, illustrated using the *Drosophila albomicans* assembly ACVV01 (Zhou et al. 2012). (**A**) Square-binned blob plot showing the distribution of assembly scaffolds on GC proportion and coverage a <u>(Challis et al.)</u>xes. Squares within each bin are coloured according to taxonomic annotation and scaled according to total span. Scaffolds within each bin can be selected for further investigation. (**B**) Cumulative assembly span plot showing curves for subsets of scaffolds assigned to each phylum relative to the overall assembly. (**C**) Snail plot (Challis et al., 2016) summary of assembly statistics. (**D**) BUSCO scores allow selection of all scaffolds with a BUSCO reference gene in each category. These images derive from analyses of the whole assembly. Each view updates automatically in response to any filters or selections that are applied to the dataset. This figure can be regenerated, and explored further, using the URLs given in File S1.

**Figure 2.**
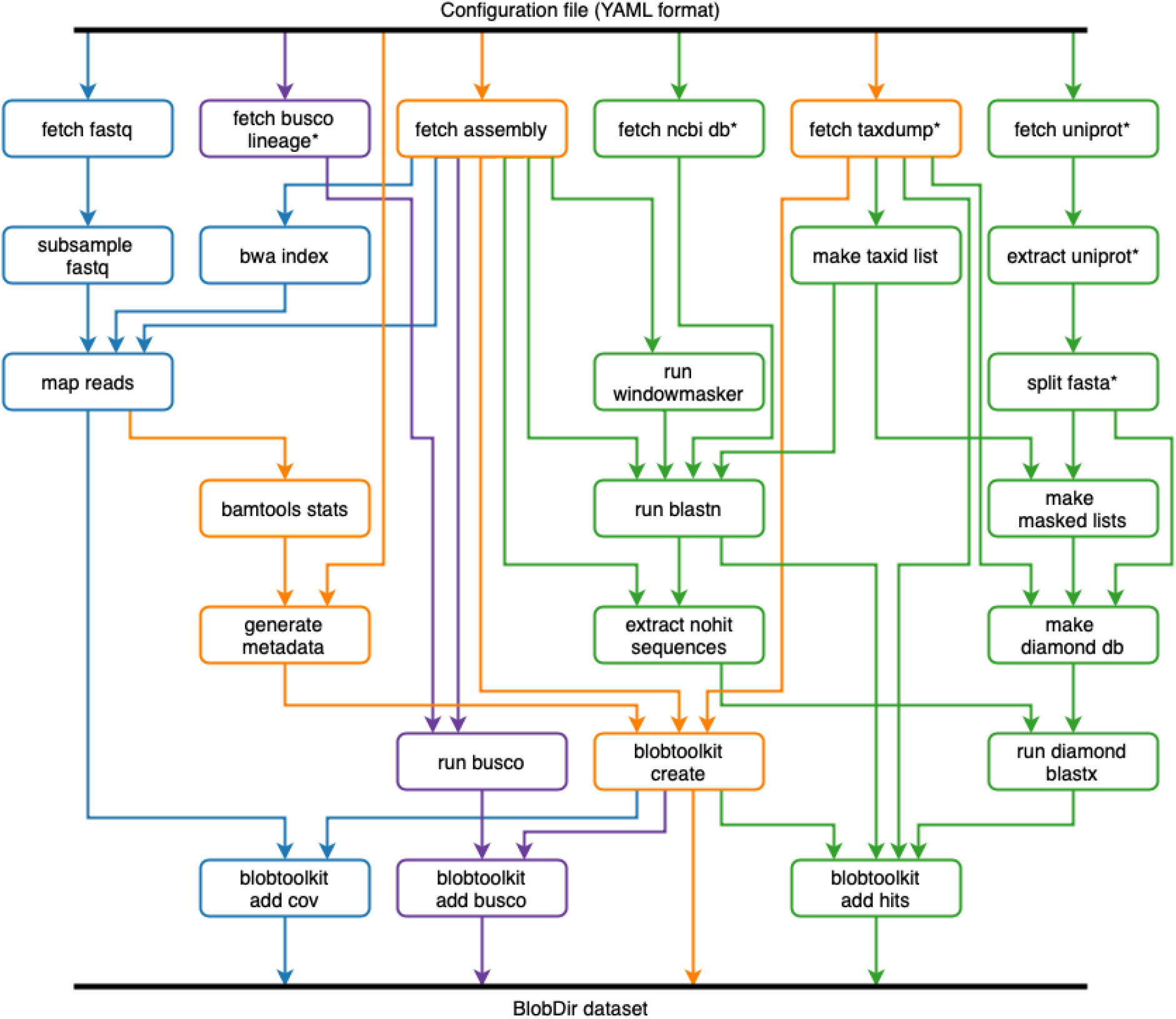
Depiction of the snakemake workflow used to analyse publicly available (INSDC-registered) eukaryotic genome assemblies. The workflow is run once for each assembly. Each box represents a Snakemake rule that may be run one or more times during workflow execution. The workflow can be logically divided into four parts: (i) creation of a minimal *BlobDir* dataset based on a single assembly with metadata derived from the configuration file and additional taxonomic annotation from the NCBI taxdump, shown in orange; (ii) addition of sequence-similarity search results based on *blastn* and Diamond *blastp* searches of the nt and refseq databases, shown in green; (iii) addition of read coverage data based on minimap2 alignment of read files linked to the assembly record (where available), shown in blue; and (iv) addition of BUSCO results based on analyses with all relevant BUSCO lineages, shown in purple. Rules marked with an asterisk are typically only run the first time the pipeline is executed as they generate local copies of relevant database files used elsewhere in the pipeline.

This workflow broadly follows the BlobTools workflow (Laetsch and Blaxter 2017), but with some changes to increase efficiency. For example, Diamond searches against UniProt are only run for contigs with no BLAST hit to the nt database, and the addition of BUSCO analyses. Query genomes are masked using *windowmasker* to reduce spurious matches to interspersed repeats. A wrapper script for *blastn* splits contigs longer than 100 kb into chunks before running BLAST, to avoid taxonomic inference for longer contigs being dependent on a single region. Since this pipeline was run on public datasets extracted from the same databases that are used to infer taxonomic affiliation, all sequences belonging to the same genus as the query assembly were excluded either before (Diamond) or during (BLAST) sequence similarity searches.

The pipeline uses Conda (https://docs.conda.io/projects/conda/en/latest/index.html) environments to load all external dependencies. These are stored as YAML-format files within the **INSDC-pipeline** repository. The *generate_metadata* step of the pipeline includes the current git commit hash in an extended version of the input configuration file so the specific versions of each program used can be determined from the *BlobDir* metadata. A record of database versions is maintained by including the date of creation in the local database directory names.

## ENA Integration

We have worked to integrate the analyses generated by BlobToolKit with the genome presentations of the European Nucleotide Archive (ENA) (Amid et al. 2019), to enhance understanding and utility of submitted data. Importantly, ENA holds both deposited raw sequence read and genome assembly data and it is possible to mine these data to discover relationships describing which read sets were used in given assemblies. At the time of analysis, of the 7,632 eukaryotic genome assemblies present within the ENA that could be associated with read sets, 585 (8%) were associated with a single run in the raw sequence data, 875 (11%) with between two to four runs, and 6,172 (81%) associated with four or more runs. None of the eukaryotic genome assemblies explicitly referenced the run(s) used to create the assembly within the relevant metadata. Values differ from those presented in Table 1, which uses only data available through the API to make associations between genome assemblies and read sets. We note that the 585 assemblies associated with a single run derived from 266 unique species, potentially permitting the identification of common contaminants in frequently-reassembled taxa. The species with the most independent assemblies were *Saccharomyces cerevisiae*, *Homo sapiens* and *Pyricularia oryzae*. These findings led to the inclusion of user documentation for the process of referencing reads during eukaryotic genome assembly submission to the ENA (https://ena-docs.readthedocs.io/en/latest/submit/assembly/genome.html#submitting-isolate-genome-assemblies). This will encourage future assemblies to be submitted with a referenced run, thereby increasing the number of assemblies for which BlobToolKit can report contamination.

A cross-reference service was set up in conjunction with in-house cloud services for the purpose of processing eukaryotic genome assemblies hosted on the ENA via BlobToolKit, as well as hosting the resulting visual and textual data. The BlobToolKit API was used to access relevant data for each assembly in coordination with Jupyter Notebooks, generating hypertext markup language (HTML) documents for assemblies with links out to associated interactive BlobToolKit Viewer analyses. Each of these documents displays the respective blob, snail and cumulative length of scaffold by phylum plots, along with assembly statistics directly from the ENA website (see, for example, https://www.ebi.ac.uk/ena/browser/view/GCA_000298335). The generation of these documents is modified autonomously based upon the data available via the API, and uploaded to GitHub Pages respectively.

## Case Studies

The following case studies highlight some of the features of BlobToolKit and the ways it may be used in assessment of published assemblies.

### Identification of common cobionts

The *Drosophila albomicans* assembly ACVV01 (GCA_000298335.1) contains 1,440 scaffolds that have greatest sequence similarity to Proteobacteria sequences in the reference databases (nt and UniProt). On a blob plot of GC proportion *vs*. coverage, many of these scaffolds are found in a distinct blob with higher GC proportion and lower coverage than the majority of the assembled scaffolds (Figure 3A and B). The difference in the distributions of the two sets is highlighted in a kite representation of the data (Figure 3B).

**Figure 3.**
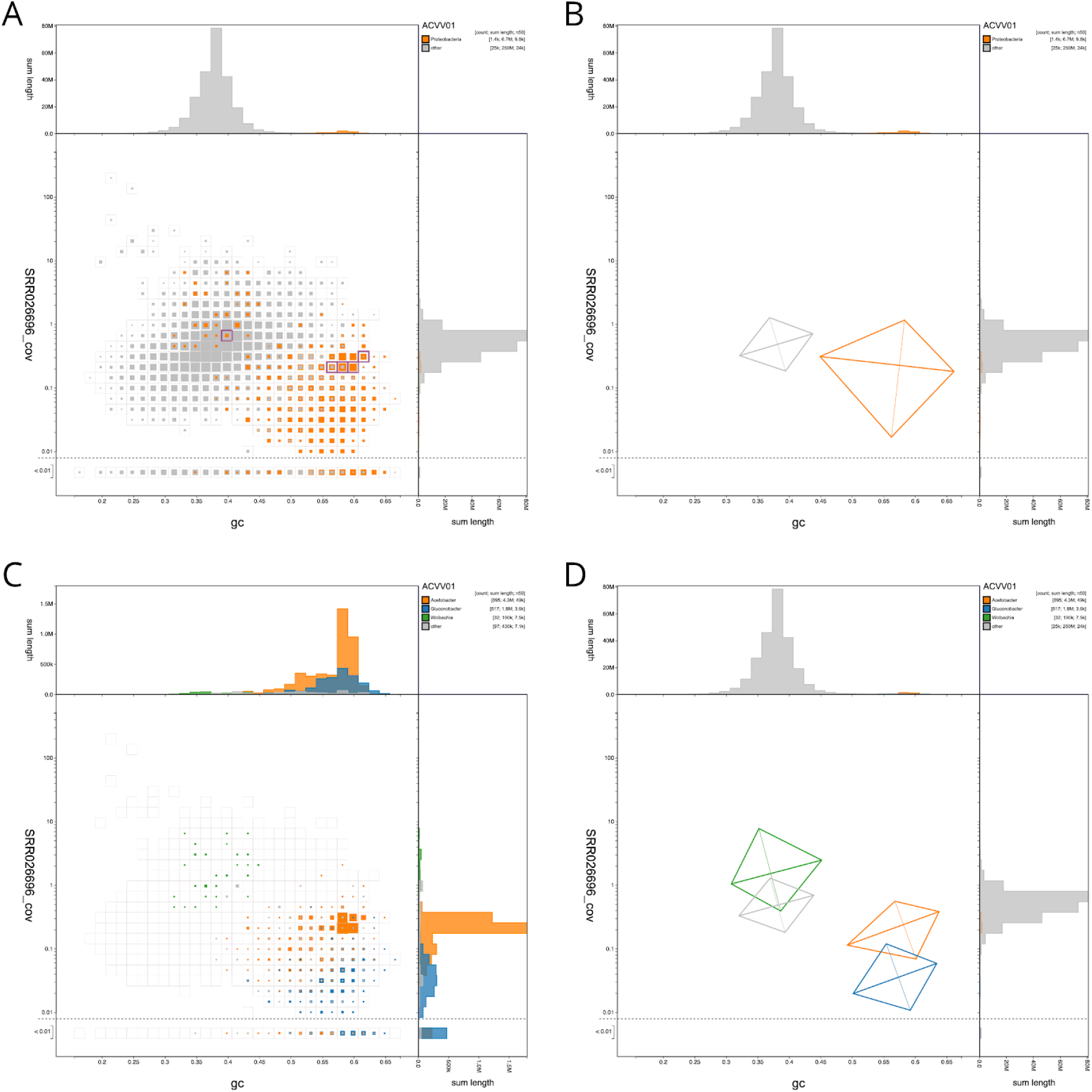
Blobplot of base coverage in read set SRR026696 against GC proportion for scaffolds in *Drosophila albomicans* assembly ACVV01. (**A** & **B**) Scaffolds are coloured by phylum with Proteobacteria highlighted in orange and all other phyla grouped together in grey. Histograms show the distribution of scaffold length sum along each axis. (**A**) Square-binned blob plot at a resolution of 30 divisions on each axis. Coloured squares within each bin are sized in proportion to the sum of individual scaffold lengths on a logarithmic scale, ranging from 867 to 40,536,114. The bins highlighted in pink contain a total of 5 scaffolds that have been annotated as Proteobacteria but that contain BUSCOs using the diptera_odb9 BUSCO set. The list of selected scaffolds is included in File S2. (**B**) A simplified representation of the distributions of scaffolds assigned to each phylum highlights the difference in GC proportion and coverage of Proteobacteria scaffolds. Each kite has a pair of lines representing two standard deviations about the mean on each axis (weighted to account for scaffold lengths) that intersect at a point representing the weighted median. They are angled according to a weighted linear regression equation to indicate the relationship between coverage and GC proportion. (**C**) Assembly filtered to exclude non-proteobacterial scaffolds. Scaffolds are coloured by genus with *Acetobacter* highlighted in orange, *Gluconobacter* shown in blue and *Wolbachia* shown in green. Coloured squares within each bin are sized in proportion to the sum of individual scaffold lengths on a square-root scale, ranging from 1,005 to 771,195. (**D**) A simplified representation of the distributions of scaffolds assigned to each genus highlights the difference in GC proportion and coverage of *Acetobacter*, *Gluconobacter* and *Wolbachia* scaffolds. This figure can be regenerated, and explored further, using the URLs given in File S1.

When analysed at higher taxonomic resolution, the scaffolds assigned to Proteobacteria derive from several distinct species. The majority of proteobacterial scaffolds (representing 4.3 Mb of 6.7 Mb) are assigned to *Acetobacter*, and there are 1.8 Mb of scaffolds assigned to *Gluconobacter* (Figure 3C). The *Gluconobacter* scaffolds have a lower coverage than the *Acetobacter* scaffolds, and thus the assembly is, as expected, less complete. *Acetobacter* and *Gluconobacter* species are common cobionts of *Drosophila* (Crotti et al. 2010) and usually have genomes of 3-4 Mb. A third group of scaffolds is assigned to the alphaproteobacterial genus *Wolbachia* (Figure 3D). *Wolbachia* are intracellular symbionts that commonly manipulate the reproductive biology of their hosts (Werren, Baldo, and Clark 2008), and insect-infecting strains have genomes of ~1.4 Mb. However, the cumulative span of scaffolds assigned to *Wolbachia* is only 190 kb. The GC proportion and coverage of these scaffolds is more congruent with that of the bulk, *Drosophila*-assigned scaffolds. Collectively, these data suggest that the *Wolbachia*-assigned scaffolds are likely to represent nuclear insertions of *Wolbachia* fragments. Such insertions are common in insect genomes, and derive from previous colonisation of the species by this endosymbiont (Dunning-Hotopp et al. 2007).

It is notable that some of the loci identified using the diptera_odb9 BUSCO set (EOG091502LX, EOG091505EO, EOG091502SD, EOG091504TW, EOG09150B43, EOG09150529) are annotated as being present in scaffolds that have been assigned to Proteobacteria. Five of these scaffolds have GC proportions and coverages consistent with their being part of the bacterial rather than the *Drosophila* genomes. Thus the BUSCO assessment of ACVV01 is compromised by the presence in the bacteria of loci which are recognised as being members of the BUSCO dipteran reference gene set. While excluding the BUSCOs identified in the proteobacterial genomes makes a very small difference to the overall BUSCO completeness score of assembly ACVV01 (83.7% *vs* 83.9% complete; diptera_odb9; BUSCO 3.0.2), their inclusion in, for example, phylogenomic analyses would lead to erroneous inferences. Similar patterns are observed in other *Drosophila* assemblies. For example, in *Drosophila elegans* assembly AFFF02, two diptera_odb9 BUSCOs (EOG0915021D, EOG091501A1) are present on scaffolds assigned to Proteobacteria. The mis-annotated BUSCOs from proteobacterial scaffolds in ACVV01 are found within core Arthropoda scaffolds in AFFF02 and *vice versa*. This highlights the importance of determining assembly integrity and contamination before assessing quality and completeness, and before proceeding to downstream analyses.

### Visualisation of highly fragmented assemblies

*Conus consors* is a cone snail studied for its production of neurotoxins (Andreson et al. 2019). The *C. consors* assembly SDAX01 (GCA_004193615.1; see https://www.ncbi.nlm.nih.gov/genome/24193) highlights the challenges associated with visualisation of highly fragmented datasets. The 2 Gb assembly is split into 2,688,687 scaffolds with an N50 length of 1,128 bp. While the full dataset can be viewed in the BlobToolKit **Viewer**, interactive visualisation of so many contigs requires use of a device with a relatively high-specification (at least 8 GB RAM) and a browser that does not limit the amount of available RAM (e.g. Firefox). To allow such assemblies to be viewed on any device, we have set default parameters to limit the computation required.

The default, binned view (Figure 4A) ensures that the number of graphic elements that must be rendered by the browser does not increase linearly with dataset size as would be the case if each scaffold were plotted individually. This representation is sufficient to show that SDAX01 has a unimodal distribution on both the GC proportion and coverage axes. However 550,837 scaffolds with a total span of over 170 Mbp have coverage below 0.01 with the selected read set (SRR1714990). An assembly of this size is typically based on a number of sequencing runs and in this case nine short read accessions are associated with the same bioproject (PRJNA267645) as the assembly. The largest three of these read sets were mapped to the assembly, allowing comparison of coverage across libraries. For scaffolds with coverage <= 0.01 in SRR1714990, a coverage *vs*. coverage plot of the remaining two libraries (SRR1719763 and SRR1712902; Figure 4B) shows the majority of these scaffolds (433,970 scaffolds with a total span of over 136 Mb) have coverage in at least one other library. Some have no coverage in any of the three libraries. It might be prudent to consider all these contigs as questionable components of the *C. consors* genome, or artefacts due to heterozygosity or misassembly.

**Figure 4.**
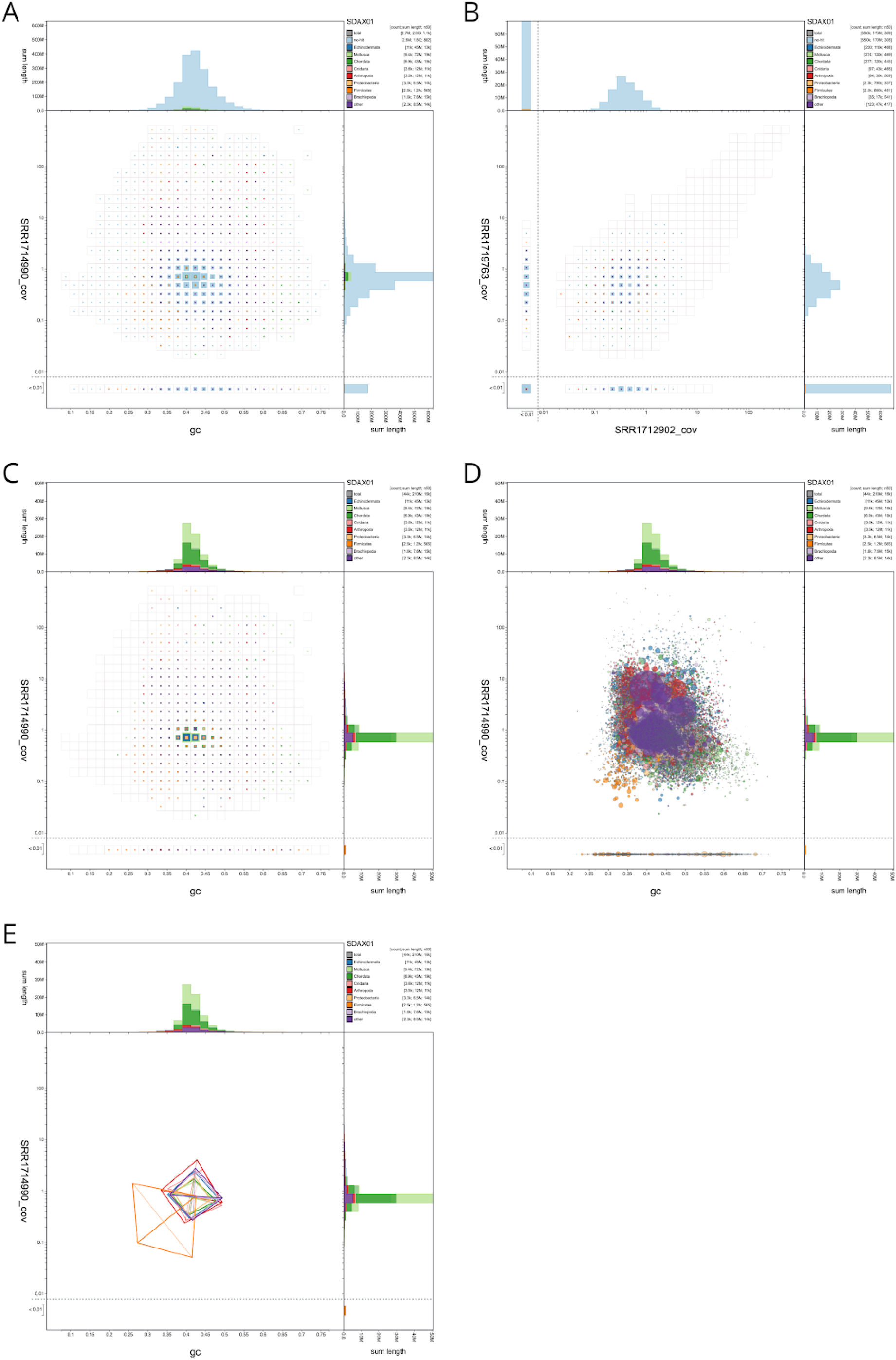
Visualisation of the highly fragmented *Conus consors* assembly SDAX01. (**A**) Binned distribution of all 2,688,687 assembly scaffolds shows unimodal distributions in GC proportion and coverage axes. The majority of scaffolds lack a taxonomic annotation (assigned to “no-hit”). (**B**) Square-binned plot of coverage in read set SRR1719763 against coverage in SRR1712902 for scaffolds with coverage <= 0.01 in read set SRR1714990. The extent of the unfiltered distribution is indicated by the empty square bins. (**C**) In the interactive browser datasets with over 1,000,000 scaffolds are presented with the “no-hit” scaffolds filtered out to reduce computation. In this case, 43,857 scaffolds are plotted in the filtered dataset. (**D**) A non-binned presentation of the same data shows the challenges of interpreting a dataset plotted as a large number of overlapping circles, even after filtering “no-hit”. (**E**) A simplified representation of the distributions of scaffolds assigned to each phylum highlights the difference in GC proportion and coverage of scaffolds assigned to Firmicute. This figure can be regenerated, and explored further, using the URLs given in File S1.

On the public BlobToolKit **Viewer** site, all datasets with over 1 million scaffolds are presented with a set of pre-generated images so users not wishing to explore beyond the default visualisations have no need to download or process the data files. In interactive mode, the same threshold is used to filter out scaffolds that lack a taxonomic annotation (those assigned to the “no-hit” category) so the default interactive view emphasises the portion of the dataset that provides most information for contaminant screening (Figure 4C). For this assembly, filtering out “no-hit” scaffolds leaves 43,857 scaffolds (1.6% of all contigs) with a total span of 209 Mb (10.2% of the total span). Below a default threshold of 100,000 scaffolds, it is computationally reasonable to plot individual scaffolds as scaled circles, even on relatively low-powered devices. However, the resulting image can be difficult to interpret as the visibility of specific features becomes dependent on plotting order with the last plotted scaffolds having greatest prominence (Figure 4D). Using a kite representation highlights a distinct distribution of Firmicute scaffolds in the *C. consors* assembly (Figure 4E) suggesting that these represent a contaminant.

### Identification of mis-annotated records in public databases

The genomes of many bird species are being generated to understand the evolution of this important group, and to explore the evolutionary genomics of particular phenotypes (Jarvis et al. 2014). While most other palaeognath birds (kiwis, ostriches, rheas and their kin) are flightless, tinamous can fly, and genomic analyses are exploring the biology of this phenotypic shift (Sackton et al. 2019). The genome assembly of the thicket tinamou, *Crypturellus cinnamomeus* (PTEZ01; GCA_003342915.1) (Sackton et al. 2019) was analysed using BlobToolKit. We noted that this assembly (total span 1.1 Gb) contained ~130 Mb of scaffolds that had coverage an order of magnitude lower than that of the main part of the assembly (Figure 5A). This blob of scaffolds also had a mean GC proportion of 0.52, contrasting with the main assembly GC proportion of 0.42. Exploring the biology of this set of scaffolds revealed several interesting features.

**Figure 5.**
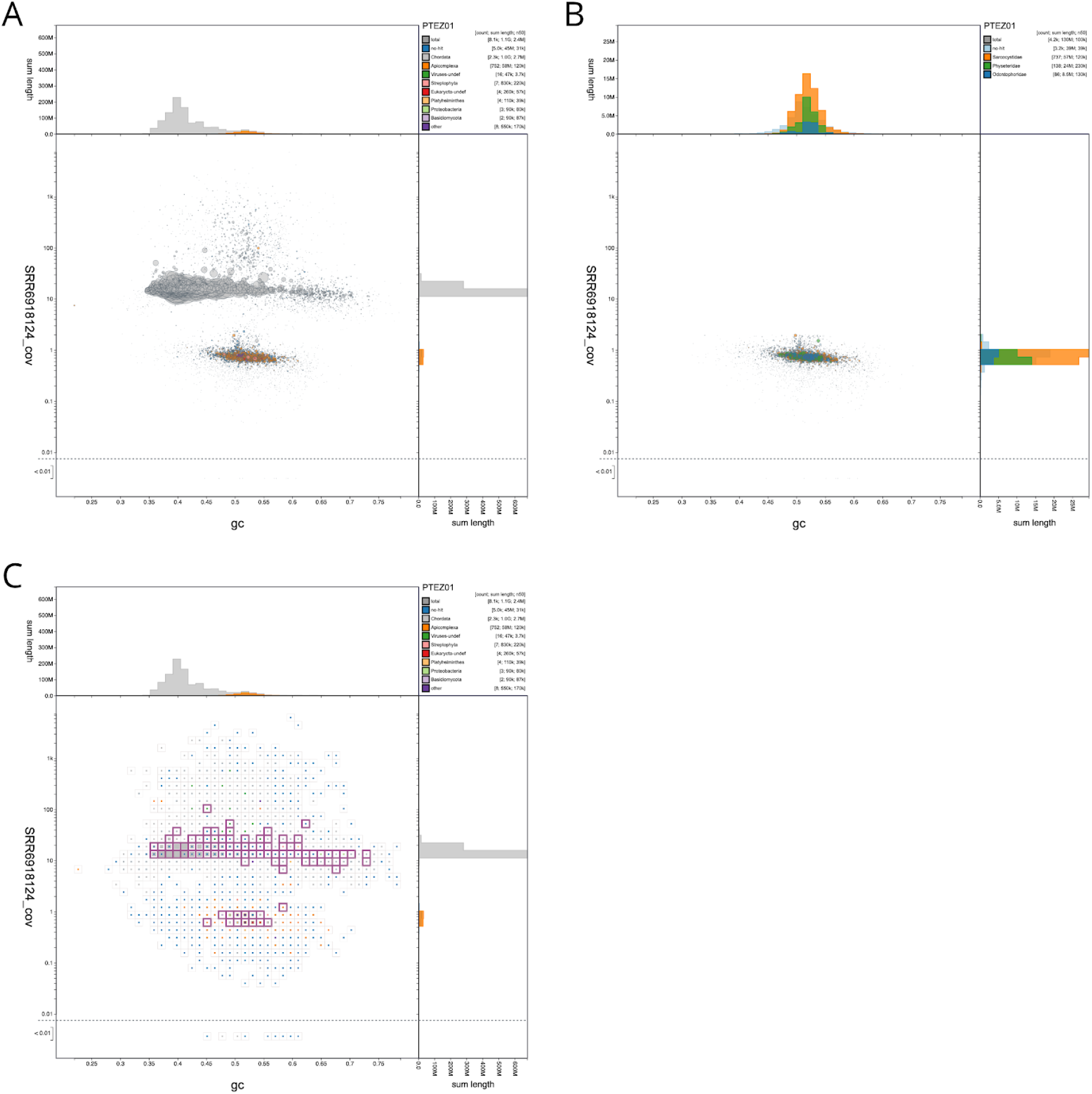
Blob plots of the *Crypturellus cinnamomeus* assembly PTEZ01 showing the presence of an apicomplexan parasite. (**A**) Circles are scaled with area proportional to scaffold length and coloured by phylum. Scaffolds assigned to the phylum Apicomplexa are coloured orange and form a distinct blob relative to the majority of Chordata-assigned scaffolds, shown in grey. (**B**) Circles are coloured by family and scaffolds assigned to families other than Physeteridae, Odontophoridae or Sarcocystidae have been filtered out. Scaffolds with coverage greater than 2 in the SRR6918124 read set have also been excluded. (**C**) A square-binned plot in which bins containing scaffolds with BUSCO annotations using any of the applicable reference gene sets are outlined in pink. The list of selected scaffolds is included in File S2. This figure can be regenerated, and explored further, using the URLs given in File S1.

Half of the span of the low-coverage scaffolds (58 Mb) was assigned to the protist group Eucoccidiorida, and more specifically had high-scoring matches to *Sarcocystis* species (Figure 5B). *Sarcocystis* are apicomplexan parasites that infect a wide range of vertebrate and non-vertebrate hosts. Sarcocystidae, which includes the important pathogens *Neospora* and Toxoplasma, have genomes that range from ~60 Mb to 127 Mb (*Sarcocystis neurona*). The other scaffolds in the low-coverage blob either had no annotation (39 Mb) or were annotated as deriving from a cetacean, *Physeter catodon* (Physeteridae; the sperm whale; 24 Mb) or a galliform bird, *Colinus virginianus* (Odontophoridae; the northern bobwhite quail; 8 Mb). While it is possible that a bird genomics laboratory might contaminate across species, the northern bobwhite genome was not sequenced by the same team that sequenced the tinamou, and contamination with sperm whale is hard to imagine. Instead, we infer that the bobwhite and sperm whale genomes are also contaminated by co-assembled genomes from *Sarcocystis*-like apicomplexans. Available *C. virginianus* and *P. catodon* assemblies were analysed with BlobToolKit to determine the presence of Apicomplexan-assigned scaffolds in these assemblies (Table 3). A total of 48 Mb of the 1.2Gb (4%) of the *C. virginianus* assembly AWGT02 (GCA_000599465.2 (Oldeschulte et al. 2017)) is inferred to be derived from an apicomplexan parasite. For *P. catodon*, the only published assembly, AWZP01 (GCA_000472045.1 (Warren et al. 2017)) is inferred to be free of contamination with sequences of apicomplexan origin. However, two more recent assemblies, including a chromosome-level assembly PGGR02 (GCA_002837175.2), which is tagged as the RefSeq (Pruitt, Tatusova, and Maglott 2005) representative genome, each contain 4.3 Mb of sequence assigned to Apicomplexa.

Thus 11% of this genome assembly appears to derive not from the target species but rather from a parasite, and sequence from this group of parasites is also present in other genome assemblies from diverse target species. This contamination of the INSDC databases with whole genomes mistakenly attributed to their host species identity means that the public commons becomes an untrustworthy substrate for discovery research. Critically, as with the *D. albomicans* example above (Figure 1), the likely *Sarcocystis-*derived scaffolds contained many BUSCO annotations (Figure 5C), and contributed 6% of the unique eukaryote BUSCO hits in the assembly (Table 2).

**Table 2.**
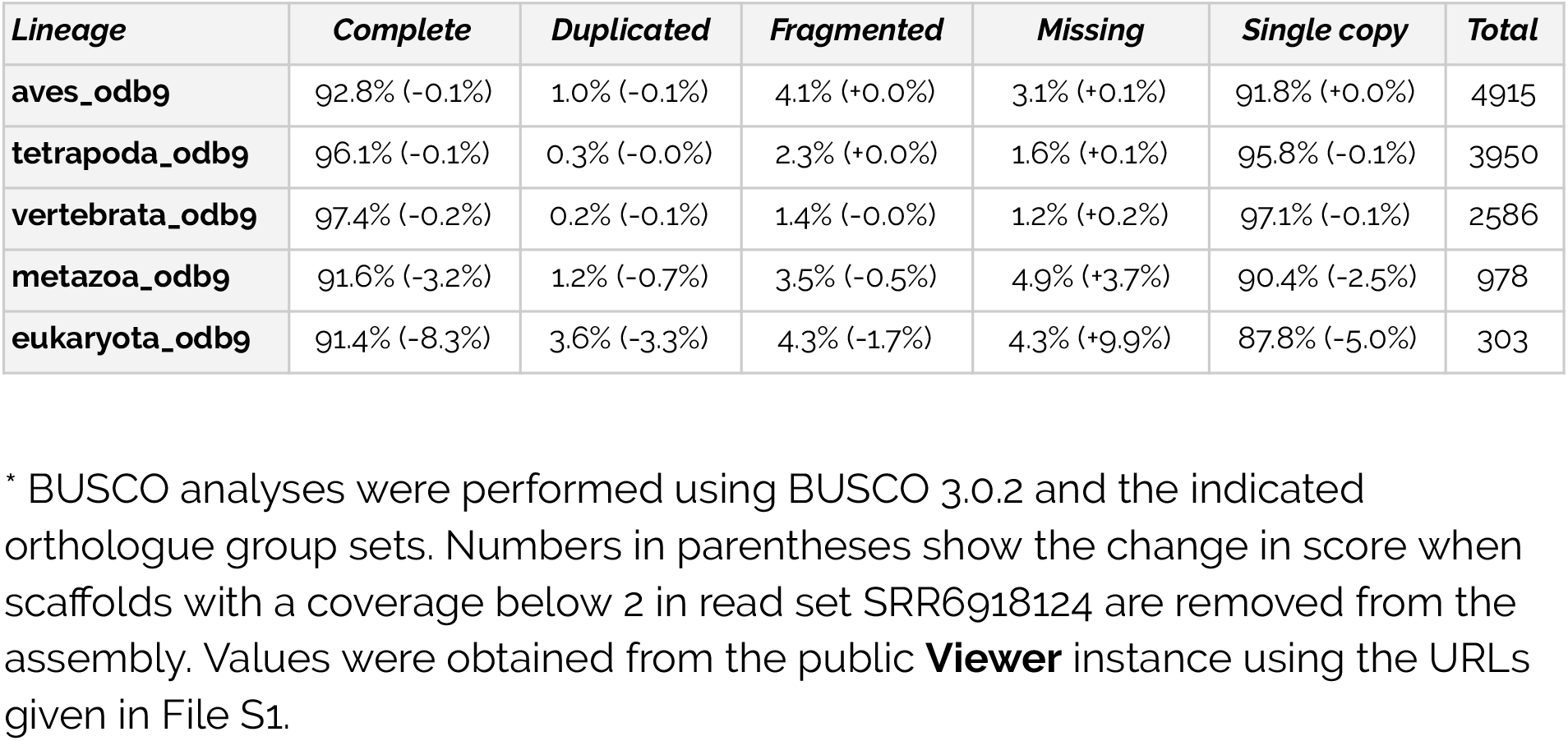
BUSCO scores* for the *Crypturellus cinnamomeus* assembly PTEZ01.

We have identified additional examples of co-sequencing of apicomplexan pathogens with target species in other taxa (Table 3). These include early assemblies of the model organisms Mus musculus and Rattus norvegicus, for which subsequent revisions have been released that have shorter span and few or no remaining apicomplexan-assigned sequences. For non-model organisms the resources available for assembly revision are considerably smaller so it is important to have the means to identify co-sequencing with pathogens and other cobionts. BlobToolKit makes evident these fascinating biological juxtapositions, and facilitates evidence-led separation of host from cobiont. Indeed this task was one of the original motivations for the development of the blob plot: to separate symbiont genomes from those of their hosts (Kumar et al. 2013).

**Table 3.** Presence of Apicomplexa-assigned sequences in selected chordate genome assemblies*.

| *Species* | *Accession* |  | *Date* | *Span (Mb)* |  |  |
| --- | --- | --- | --- | --- | --- | --- |
|  | *BlobToolKit* | *GCA* |  | *Apicomplexa* | *Chordata* | *Total* |
| *Colinus virginianus* | AWGT02 | GCA_000599465.2 | 2017<sup>a</sup> | 48.0 | 1074 | 1254 |
| *Crypturellus cinnamomeus* | PTEZ01 | GCA_003342915.1 | 2018<sup>b</sup> | 59.9 | 1017 | 1122 |
| *Mus musculus* | AAHY01 | GCA_000002165.1 | 2009<sup>c</sup> | 188.9 | 2738 | 3251 |
| *Mus musculus* | LXEJ02 | GCA_003774525.2 | 2018<sup>d</sup> | 0.0 | 2687 | 2801 |
| *Physeter catodon* | AWZP01 | GCA_000472045.1 | 2013<sup>e</sup> | 0.0 | 2279 | 2280 |
| *Physeter catodon* | UEMC01 | GCA_900411695.1 | 2018<sup>f</sup> | 4.3 | 2472 | 2512 |
| *Physeter catodon* | PGGR02 | GCA_002837175.2 | 2019<sup>g</sup> | 4.3 | 2472 | 2512 |
| *Piliocolobus tephrosceles* | PDMG02 | GCA_002776525.2 | 2018<sup>h</sup> | 33.3 | 2976 | 3038 |
| *Rattus norvegicus* | AAHX01 | GCA_000002265.1 | 2006<sup>i</sup> | 15.4 | 2699 | 2932 |
| *Rattus norvegicus* | AABR07 | GCA_000001895.4 | 2014<sup>j</sup> | 0.2 | 2869 | 2870 |

\* The data for this table can be obtained using the URL given in File S1
<sup>a</sup> (Oldeschulte et al. 2017)
<sup>b</sup> (Sackton et al. 2019)
<sup>c</sup> (Mural et al. 2002)
<sup>d</sup> Most recent non-chromosomal assembly (see
https://www.ncbi.nlm.nih.gov/assembly/GCA_003774525.2)
<sup>e</sup> (Warren et al. 2017)
<sup>f</sup> (see https://www.ncbi.nlm.nih.gov/assembly/GCA_900411695.1)
<sup>g</sup> (see https://www.ncbi.nlm.nih.gov/assembly/GCA_000472045.1)
<sup>h</sup> (see https://www.ncbi.nlm.nih.gov/assembly/GCA_000472045.1)
<sup>i</sup> (Florea et al. 2005)
<sup>j</sup> (Gibbs et al. 2004)

## Discussion

BlobToolKit is a significant extension of the approach launched in BlobTools. In particular, by permitting user interaction with the rich data associated with each contig in the **Viewer** mode, BlobToolKit can enhance discovery of novel biology. The addition of real-time interaction addresses a criticism of the approach, relative to cluster-based methods such as Anvi’o (Eren et al. 2015), that it limits the amount of supporting data that can be included (Delmont and Eren 2016). We envisage three main uses for BlobToolKit. The first is in the research laboratory aiming to sequence for the first time the genome of a new species. BlobToolKit can be used during the assembly process, to filter contaminants and cobionts, and to explore issues such as haploid *versus* diploid contigs, and patterns of coverage in different sequence read datasets (for example, comparing male and female read sets in heterogametic organisms). As part of an assembly workflow, BlobToolKit should ensure better quality assemblies with higher biological credibility.

The second use is in publication and visualisation of published assemblies. The BlobToolKit **Viewer** generates publication quality images that are fully reproducible via the embedding of control parameters in the URL. These images should, we believe, become standard in reporting genome assemblies, and thus enhance the ease of assessment of assembly quality. We have worked to embed BlobToolKit views into the presentation of genome assemblies at the ENA for just this reason and believe that we have demonstrated that collaboration between tools developers and public databases is important in refining best practice in data publication. Journals may generate (or request that authors supply) BlobToolKit assessments of new assemblies submitted for publication, to aid review and speed publication of high quality data.

The third is in comparative and evolutionary genomics. With ongoing improvements in sequencing technologies and assembly software, genome assemblies are improving in quality and contiguity. Amongst other players, the Earth Biogenome Project (Lewin et al. 2018), 10K Vertebrate Genome Project (Genome 10K Community of Scientists 2009) and Tree of Life project (https://www.sanger.ac.uk/science/programmes/tree-of-life) collectively aim to generate chromosomally-contiguous reference genomes for (in the first instance) all known families of Eukaryota. BlobToolKit protocols can be used to explore these genomes for evidence of past horizontal gene transfer, for the presence of symbionts and parasites, and to explore chromosomal patterns of gene expression.

The difficulty we experienced in associating raw sequence read sets with submitted assemblies has led ENA to include a more apparent and thorough explanation of the benefits of and process for referencing reads during eukaryotic genome assembly submission to the repository. We advocate the practice of assembly submission along with associated reads to INSDC to enable downstream analysis and assembly contamination detection.

We aim to complete analysis of all public genomes in INSDC and post them to the BlobToolKit **Viewer** website at https://blobtoolkit.genomehubs.org/view in the near future, and then maintain currency with the flow of new genomes. The toolkit is under active development (see https://github.com/blobtoolkit) and we welcome feature requests and collaborations to expand and improve its capabilities.

## Data Availability

All code is available on Github and release versions have been deposited in the Zenodo open access repository:

**BlobTools2** v2.1:

https://github.com/blobtoolkit/blobtools2

https://doi.org/10.5281/zenodo.3531583

**INSDC-pipeline** v1.0:

https://github.com/blobtoolkit/insdc-pipeline

https://doi.org/10.5281/zenodo.3533168

**Specification** v1.0:

https://github.com/blobtoolkit/specification

https://doi.org/10.5281/zenodo.3531846

**Viewer** v1.0:

https://github.com/blobtoolkit/viewer

https://doi.org/10.5281/zenodo.3533128

All processed datasets referred to in this manuscript can be viewed interactively on the public instance of the **Viewer** at https://blobtoolkit.genomehubs.org/view, accessed programmatically through the **Viewer** API (see https://blobtoolkit.genomehubs.org/api-docs/) or downloaded from https://blobtoolkit.genomehubs.org/download.

Supplementary Files S1-S3 have been deposited in Figshare.

## Acknowledgements

BlobToolKit is based on Blobology by Sujai Kumar and BlobTools by Dominik Laetsch and we thank both, and other colleagues in the Blaxter lab, for their comments and criticisms. This work was funded by a BBSRC Bioinformatics and Biological Resources award BB/P024238/1.

